# Rapid assessment of SARS-CoV-2 evolved variants using virus-like particles

**DOI:** 10.1101/2021.08.05.455082

**Authors:** Abdullah M. Syed, Taha Y. Taha, Mir M. Khalid, Takako Tabata, Irene P. Chen, Bharath Sreekumar, Pei-Yi Chen, Jennifer M. Hayashi, Katarzyna M. Soczek, Melanie Ott, Jennifer A. Doudna

## Abstract

Newly evolved SARS-CoV-2 variants are driving ongoing outbreaks of COVID-19 around the world. Efforts to determine why these viral variants have improved fitness are limited to mutations in the viral spike (S) protein and viral entry steps using non-SARS-CoV-2 viral particles engineered to display S. Here we show that SARS-CoV-2 virus-like particles can package and deliver exogenous transcripts, enabling analysis of mutations within all structural proteins and rapid dissection of multiple steps in the viral life cycle. Identification of an RNA packaging sequence was critical for engineered transcripts to assemble together with SARS-CoV-2 structural proteins S, nucleocapsid (N), membrane (M) and envelope (E) into non-replicative SARS-CoV-2 virus-like particles (SC2-VLPs) that deliver these transcripts to ACE2- and TMPRSS2-expressing cells. Using SC2-VLPs, we tested the effect of 30 individual mutations within the S and N proteins on particle assembly and entry. While S mutations unexpectedly did not affect these steps, SC2-VLPs bearing any one of four N mutations found universally in more-transmissible viral variants (P199L, S202R, R203M and R203K) showed increased particle production and up to 10-fold more reporter transcript expression in receiver cells. Our study provides a platform for rapid testing of viral variants outside a biosafety level 3 setting and identifies viral N mutations and viral particle assembly as mechanisms to explain the increased spread of current viral variants, including Delta (N:R203M).

**One-Sentence Summary:** R203M substitution within SARS-CoV-2 N, found in delta variant, improves RNA packaging into virus-like particles by 10-fold.

## Main Text

The COVID-19 pandemic is a leading cause of death globally due to the ongoing emergence of SARS-CoV-2 variants with increased transmissibility. Understanding the molecular determinants of enhanced infectivity is central to vaccine and therapeutic development, but research is hindered because SARS-CoV-2 can only be handled safely in a biosafety level 3 (BSL-3) lab. Furthermore, generating mutant infectious clones of SARS-CoV-2 is technically challenging and risks generation of viruses with increased virulence (*1*–*5*). Current studies employ spike (S) pseudotyped lentivirus systems for rapid evaluation of S-mediated ACE2 receptor binding and cell entry (*6*, *7*). However, most mutations observed in circulating variants occur outside the S gene and are thus inaccessible by this approach (*8*).

All SARS-CoV-2 variants of interest or concern defined by the WHO contain at least one mutation with >50% penetrance within seven amino acids (N:199-205) in the nucleocapsid (N) protein, which is required for replication and RNA binding, packaging, stabilization and release (*8*). Despite its functional importance and emergence as a mutational hotspot, the N protein has not been widely studied because of the absence of simple and safe cell-based assays. Biochemical analysis of N has also proven difficult because of its instability and propensity to assemble or phase-separate and to bind RNA non-specifically (*9*–*11*). To investigate N function, effects of mutations and other aspects of SARS-CoV-2 biology, we set out to develop a SARS-CoV-2-based packaging system to deliver exogenous RNA transcripts into cells using virus-like particles (VLPs).

We reasoned that a process mimicking viral assembly to package and deliver reporter transcripts would simplify the analysis of successful virus production, budding and entry. Previous studies have shown that co-expression of only the structural proteins of coronaviruses within human cells generates VLPs containing all four structural proteins (*12*–*17*). VLPs generated using this method appear to have similar morphology to infectious viruses and have previously been proposed as vaccine candidates (*18*). A key requirement for such VLPs to deliver reporter transcripts into cells is the packaging signal necessary for RNA recognition. During natural viral assembly, the N protein is thought to recognize an RNA structure that overlaps a coding sequence within ORF1ab, enabling the full viral genome that contains this sequence to be packaged to the exclusion of viral subgenomic and host transcripts (*19*). The identification of the SARS-CoV-2 RNA packaging signal is required to create SARS-CoV-2 VLPs (SC2-VLPs) that package engineered transcripts by this mechanism.

Based on the reported packaging sequences for related viruses including Murine Hepatitis Virus and SARS-CoV-1, we hypothesized that the SARS-CoV-2 packaging signal might reside within genomic fragment “T20” (nt 20080-22222) encoding non-structural protein 15 (nsp15) and nsp16 (Fig. 1A) (*16*, *19*–*21*). We designed a transfer plasmid encoding a luciferase transcript containing T20 within its 3’ untranslated region (UTR) and tested for SC2-VLP production by co-transfecting the transfer plasmid into packaging cells (HEK293T) along with plasmids encoding the virus structural proteins (Fig. 1B). Supernatant secreted from these packaging cells was filtered and incubated with receiver 293T cells co-expressing SARS-CoV-2 entry factors ACE2 and TMPRSS2 (Fig. 1B). We observed luciferase expression in receiver cells only in the presence of all four structural proteins (S, M, N, E) as well as the T20-containing reporter transcript (Fig. 1C). Substituting any one of the structural proteins or the luciferase-T20 transcript with a luciferase-only transcript decreased luminescence in receiver cells by >200-fold and 63-fold respectively (Fig. 1C). We also conducted this experiment using Vero E6-TMPRSS2 cells that endogenously express ACE2 and once again observed robust luciferase expression only in the presence of all five components (fig. S1A).

**Figure 1:**
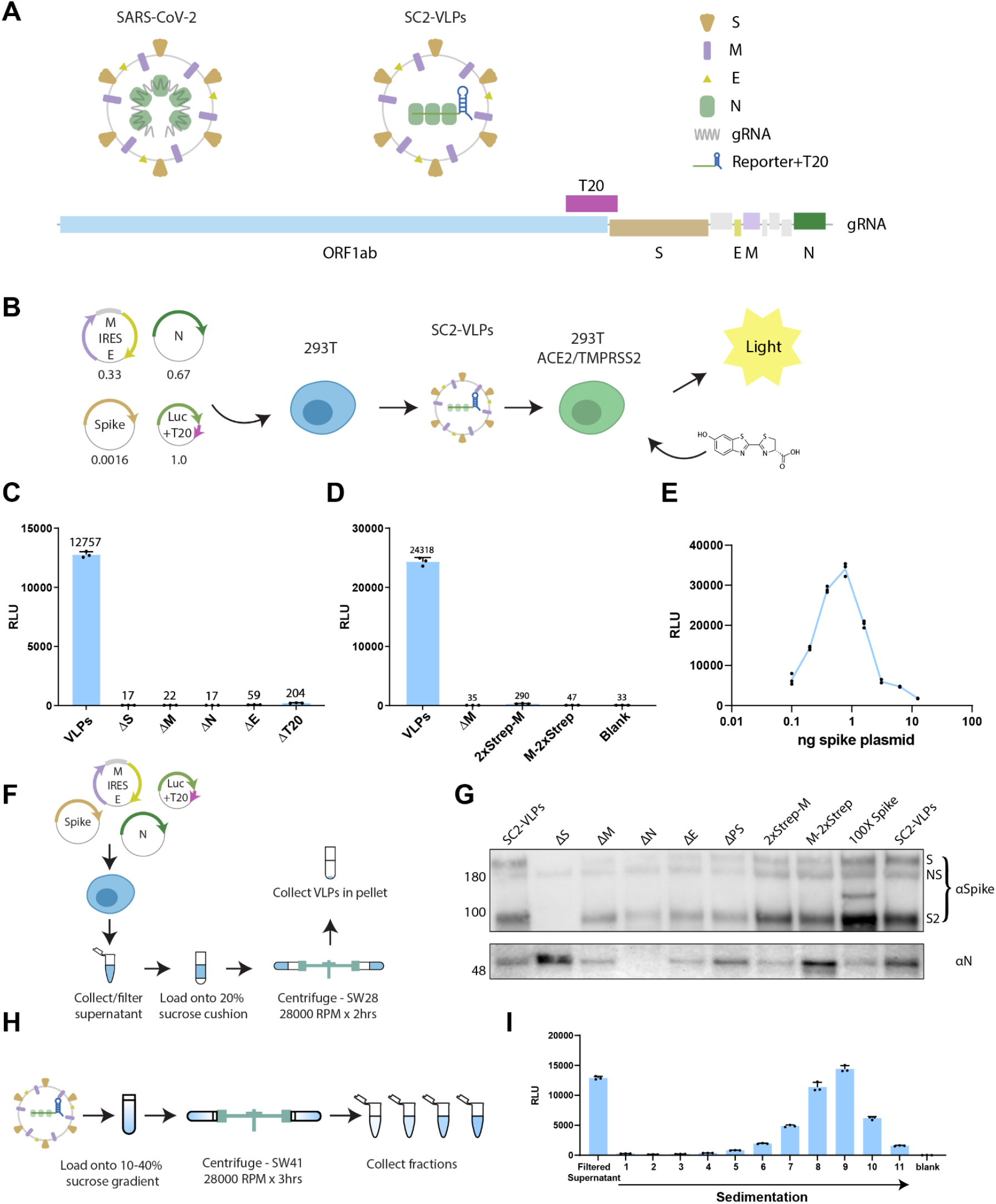
Design and characterization of SC2-VLPs. A) Schematic of SARS-CoV-2 virus and vector design. B) Process flow for generating and detecting luciferase encoding CoV-2 VLPs. Numbers below plasmid maps indicate ratios used for transfection. C) Induced luciferase expression measured in receiver cells (293T overexpressing ACE2 and TMPRSS2) from “Standard” CoV-2 VLPs containing S, M, N, E and luciferase-T20 transcript as well as VLPs lacking one of the components. D) N- or C-terminal strep-tag on membrane protein abrogates vector induced luciferase expression. E) Optimal luciferase expression requires a narrow range of spike plasmid concentrations corresponding to ~1ng of plasmid in a 24-well. F) Schematic for purification of virus-like particles (VLPs) including CoV-2 VLPs. G) Western blot showing spike and N in pellets purified from standard SC2-VLPs and conditions that did not induce luciferase expression in receiver cells. H) Schematic for sucrose gradient for separating CoV-2 VLPs. I) Induced luciferase expression from sucrose gradient fractions of CoV-2 VLPs.

Our approach required two key modifications compared to previous work on SARS-CoV-2 VLPs. First, although affinity sequence tags on N were tolerated, untagged native M protein was required for SC2-VLP-mediated reporter gene expression (Fig. 1D, S1B; tags on S and E were not tested). Second, luciferase expression in receiver cells was most efficient within a narrow range and surprisingly low ratio of S expression plasmid relative to the other plasmids (Fig. 1E). These findings contrast with detection of N and S proteins within pelleted material (Fig. 1F, G), suggesting that particles produced under less stringent conditions are not competent for delivering RNA. This may explain why exogenous RNA delivery has not been observed previously for SARS-CoV-2 VLPs.

Further analysis showed that SARS-CoV-2 VLPs (SC2-VLPs) are stable against ribonuclease A, resistant to freeze-thaw treatment (fig. S2A) and can be concentrated by precipitation, ultrafiltration and ultracentrifugation through a 20% sucrose cushion (fig. S2B). Analysis of SC2-VLPs fractionated using 10-40% sucrose gradient ultracentrifugation showed that large dense particles are responsible for inducing luciferase expression (Fig. 1H, I). These data support the conclusion that SC2-VLPs are formed under our experimental conditions and deliver selectively packaged transcripts by receptor-mediated cell entry.

Next, we used SC2-VLPs to locate more accurately the SARS-CoV-2 RNA packaging signal. We generated a library of 28 2-kB overlapping tiled segments (T1-T28) from the SARS-CoV-2 genome and inserted these fragments individually into a luciferase-encoding plasmid (Fig. 2A). SC2-VLPs generated using luciferase-encoding plasmids that included any region of ORF1ab from SARS-CoV-2 produced luminescence detectable in this assay, suggesting that packaging does not rely entirely on one contiguous RNA sequence (Fig. 2B, C). Furthermore, luciferase-encoding plasmids that included fragments T24-28 resulted in lower luciferase expression (Fig. 2B, C), consistent with natural exclusion of subgenomic viral transcripts containing these sequences to avoid generation of replication-defective virus particles. Overall, packaging was most efficient using T20 (nt 20080 - 22222) located near the 3’ end of ORF1ab (Fig. 2B, C) and partially but not completely overlapping with PS580 (19785-20348), which was predicted to be the packaging signal for SARS-CoV-1 based on structural similarity to known coronavirus packaging signals (*16*).

**Figure 2:**
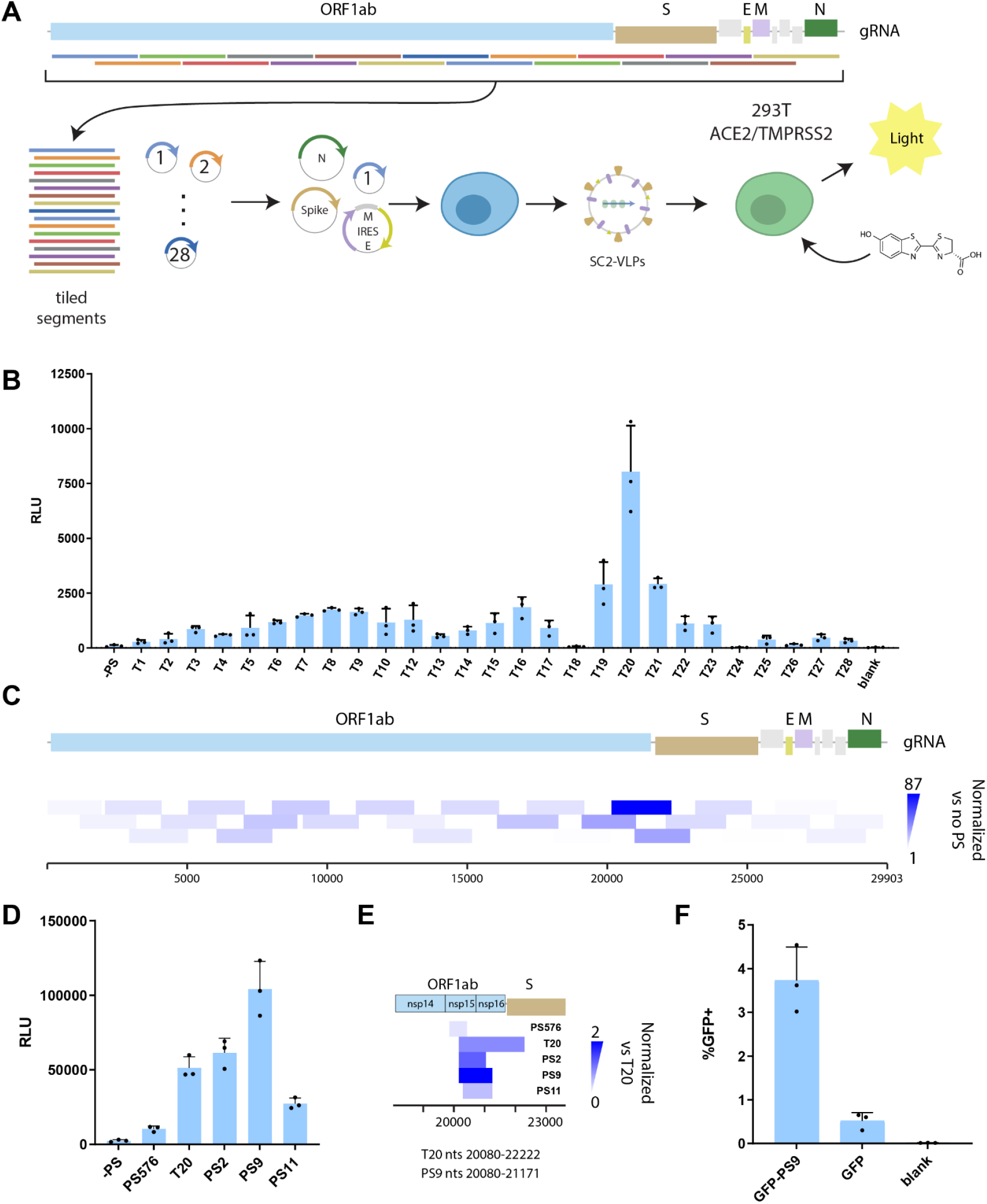
Location of the SARS-CoV-2 packaging signal. A) Arrayed screen for determining the location of the packaging signal using SC2-VLPs. 2kB tiled segments of the genome are cloned into the 3’UTR of the luciferase plasmid. B) Induced luciferase expression in receiver cells by SC2- VLPs containing different tiles from the SARS-CoV-2 genome. C) Heatmap visualization of the data from (B) showing the locations of tiled segments relative to the SARS-CoV-2 genome. Color intensity indicates luminescence of receiver cells for each tile normalized to expression for luciferase plasmid containing no insert. D) Smaller segments of the genome used to further narrow down the location of the packaging signal. E) Heatmap visualization of the data from (D). F) Flow cytometry analysis of GFP expression 293T ACE2/TMPRSS2 cells incubated with SC2-VLPs encoding GFP-PS9, GFP (no packaging signal) or no VLPs.

To further define the packaging sequence, we tested truncations and additions to T20, as well as PS580 from SARS-CoV-1. We found that PS580 resulted in lower luciferase expression compared to T20 (Fig. 2D, E; fig. S3A, B). Unexpectedly, the highest luciferase expression level resulted from SC2-VLPs encoding the nucleotide sequence 20080-21171 (termed PS9), and further truncations of this sequence reduced expression (Fig. 2D, E; fig. S3A, B). We also generated VLPs encoding GFP and found that they induced GFP expression in receiver cells in the presence of PS9 (Fig. 2F). These data suggest that PS9 (nt 20080-21171) is a *cis*-acting element that enhances RNA packaging in the presence of SARS-CoV-2 structural proteins, although our data do not exclude the presence of other *cis*-acting sequences within the SARS-CoV-2 genome.

SC2-VLPs provide a new and more physiological model compared to pseudotyped viruses for testing mutations in all four viral structural proteins (S, E, M, N) for effects on assembly, packaging and cell entry. We generated SC2-VLPs with 15 S mutations including four with the combined S mutations found in the Alpha, Beta, Gamma and Epsilon variants and surprisingly did not observe improved luciferase expression from SC2-VLPs due to any of these mutations (Fig. 3A-C). Since nearly all circulating variants contain the S:D614G mutation, we compared all mutants to the ancestral S protein modified to include G614 (termed WT+D614G). Minor changes in S expression between mutants may be a confounding factor since SC2-VLPs mediate luciferase expression optimally in a narrow range of S expression. Over a range of 6.25 ng to 50 pg per well of S-encoding plasmid (total 1 μg of DNA used in each condition), none of the tested S mutations produced >2-fold improvement in luciferase expression (Fig. 3D; fig. S4); slightly improved luciferase expression occurred with the S sequence derived from the Alpha variant (B.1.1.7) and S containing the mutation N501Y within the receptor binding domain. This finding contrasts with prior results using S-pseudotyped lentiviruses, where enhanced entry was reported for some S mutations including S:N501Y (*22*, *23*). However, S mutations tested in the context of SARS-CoV-2 infectious clones have shown mixed effects, suggesting complex or indirect connections between S and infectivity (*24*, *25*). To determine whether SC2-VLPs can accurately assess cell entry mediated by S mutants will require additional comparisons with infectious clones, but due to the lack of observed effects we instead examined mutations in N.

**Figure 3:**
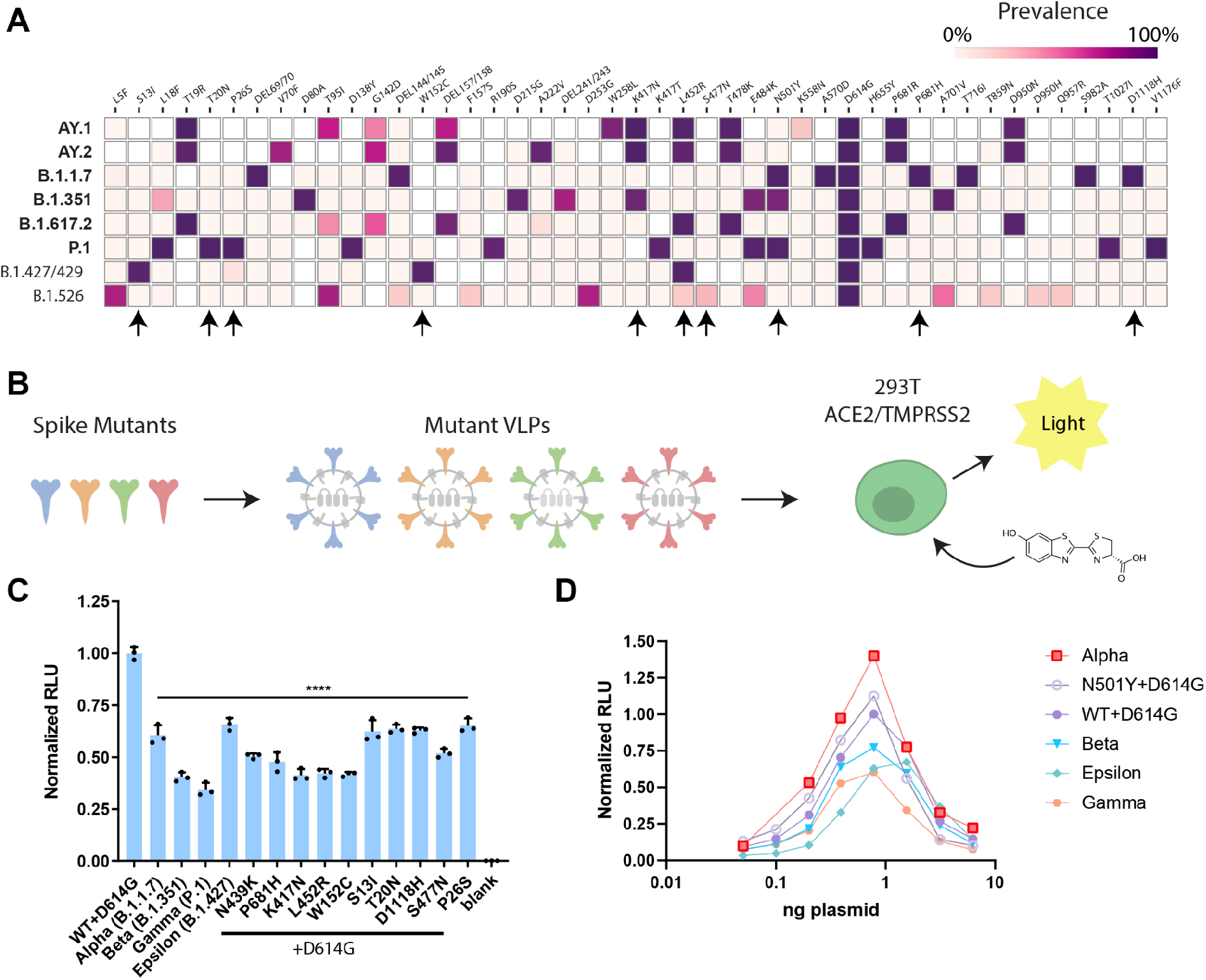
Effect of amino acid changes in the spike protein on SC2-VLP induced luminescence. A) Heatmap of observed mutations within the spike protein as of July, 2021. Each row corresponds to a variant of concern (bold) or variant of interest (regular) shown on left and each column indicates observed mutations shown at top. Colors indicate prevalence of each mutation and arrows at bottom indicate the mutations that were tested. B) Schematic for cloning and testing each variant using SC2-VLPs. C) Initial screen of 15 spike mutants compared to a reference ancestral spike containing the D614G mutation. D) Spike mutants tested at a range of plasmid dilutions with all other plasmid maintained at the same concentration.

We tested whether N mutations found in circulating SARS-CoV-2 variants result in improved viral particle assembly, RNA delivery and/or reporter gene expression using SC2-VLPs. Interestingly, half of the amino acid changes observed within N in circulating SARS-CoV-2 variants occur within a seven amino acid region (aa199-205) of the central disordered region (termed the “linker” region, Fig. 4A, B). We tested 15 N protein mutations including two combinations corresponding to the Alpha and Gamma variants since they both contain the co-occurring R203K/G204R mutations (Fig. 4B, C). The Alpha and Gamma variant N improved luciferase expression in receiver cells by 7.5- and 4.2-fold respectively relative to the ancestral Wuhan Hu-1 N-protein (Fig. 4D). In addition, four single amino acid changes improved luciferase expression: P199L, S202R, R203K and R203M. Two of these amino acid changes do not change the overall charge (P199L, R203K), one results in a more positive charge (S202R,) and one results in a more negative charge (R203M), suggesting that the improvement in luciferase expression is not likely due to simple electrostatics. Western blotting revealed no correlation between N protein expression levels and luciferase induction, suggesting that these N mutations enhance luciferase induction through a different mechanism (fig. S5).

**Figure 4:**
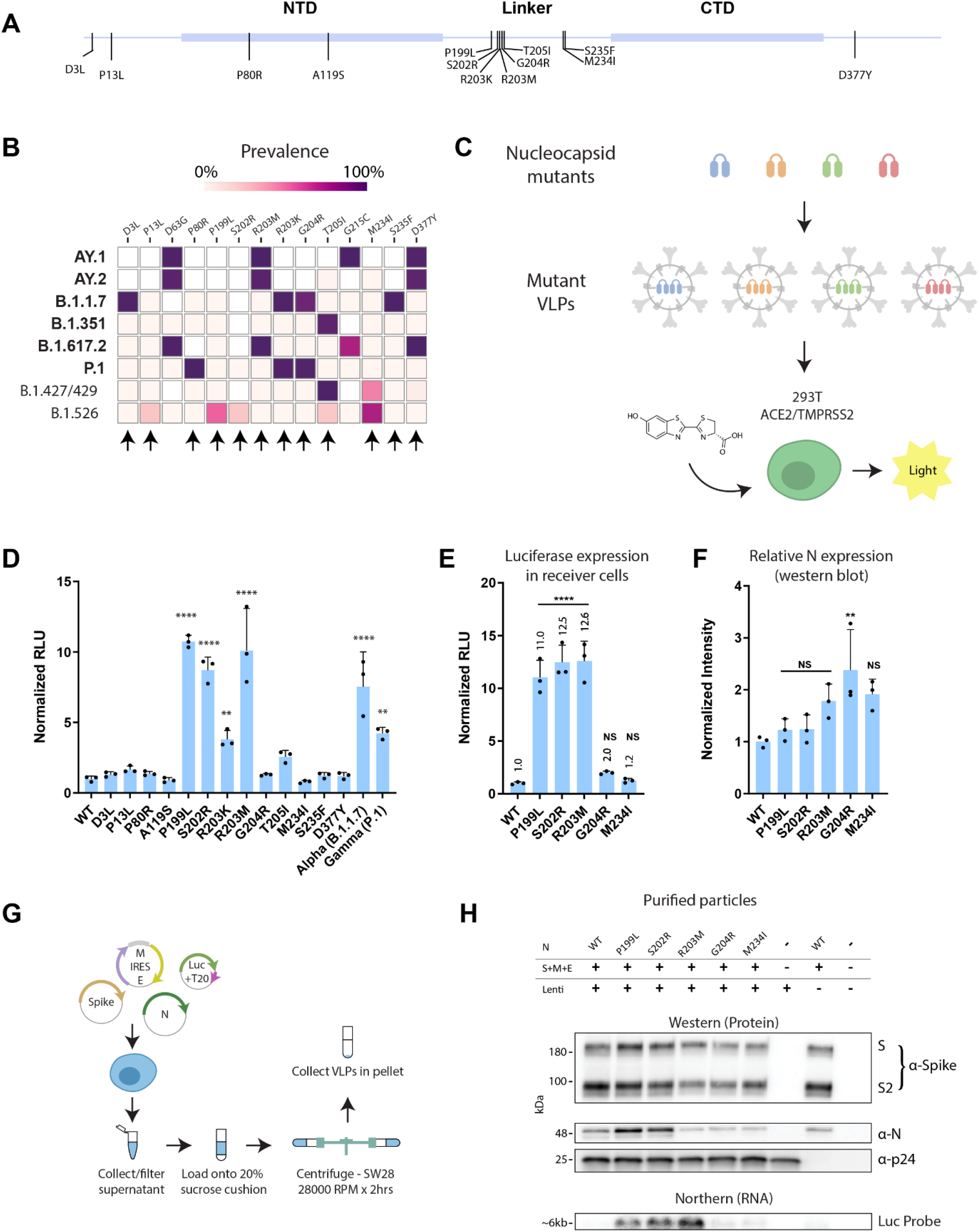
Effect of amino acid changes in the N protein on SC2-VLP induced luminescence. A) Map of SARS-CoV-2 N showing the locations of observed mutations. B) Heatmap of observed mutations within the N protein as of July, 2021. Each row corresponds to a variant of concern (bold) or variant of interest (regular) shown on left and each column indicates observed mutations shown at top. Colors indicate prevalence of each mutation and arrows indicate mutations that were tested. C) Schematic of the process flow for screening N mutations using SC2-VLPs. D) Initial screen of 15 N mutants compared to the reference Wuhan Hu-1 N sequence (WT). E) Six N mutants re-tested for luciferase expression after preparation in a larger batch. F) Relative N-expression in packaging cells normalized to WT using GAPDH as a loading control. G) Schematic for isolating purified VLPs for western and northern blot. H) Western blot (protein) and Northern blot (RNA) of isolated VLPs generated from the six N mutants as well as controls and blanks. 1mL of a batch of lentivirus was added to each sample before ultracentrifugation to allow p24 to be used as a loading control. Anti-N antibody (abcam, ab273434) binds to CTD of N which does not contain any of the mutations tested.

Further analysis of six of these N variants was conducted to determine whether these mutations affect SC2-VLP assembly efficiency, RNA packaging, or RNA uncoating prior to expression. We chose the three mutants that demonstrated ~10-fold improved luciferase expression (P199L, S202R, R203M) and two mutants that did not increase luciferase expression significantly in the preliminary screen (G204R, M234I) and compared these to the wild type (Fig. 4E). N protein expression levels in packaging cells again did not correlate with luciferase expression in receiver cells transduced with SC2-VLPs bearing these mutations. We noticed that G204R showed increased N expression in packaging cells but this did not result in a statistically significant increase in luciferase production in receiver cells (Fig. 4F). We then purified SC2-VLPs containing each N mutation and found that those containing P199L and S202R had increased levels of S, N and RNA while R203M showed increased RNA only compared to the mutants that did not demonstrate enhanced luciferase induction (Fig. 4G, H). These results suggest that mutations within the N linker domain improve the assembly of SC2-VLPs, leading either to greater overall VLP production, a larger fraction of VLPs that contain RNA or higher RNA content per particle. In either case, these results suggest a previously unanticipated explanation for the increased fitness and spread of SARS-CoV-2 variants of concern.

Overall, we present a strategy for rapidly generating and measuring SARS-CoV-2 VLPs that package and deliver exogenous RNA. This approach allows examination of viral assembly, budding, stability, maturation, entry and genome uncoating involving all of the viral structural proteins (S, E, M, N) without generating replication-competent virus. Such a strategy is useful not only for dissecting the molecular virology of SARS-CoV-2 but also for future development and screening of therapeutics targeting assembly, budding, maturation and entry. This strategy is ideally suited for the development of new antivirals targeting SARS-CoV-2 as it is highly sensitive, quantitative and scalable to high-throughput workflows. Our identification of an RNA sequence within the SARS-CoV-2 genome capable of triggering packaging of exogenous transcripts may enable the engineering of SARS-CoV-2 vaccines or therapeutics. It may be possible to introduce silent mutations within the packaging signal sequence to generate weakened strains of SARS-CoV-2 for use as an infectious vaccine or to generate defective genomes that package more efficiently than the original virus for use as a therapeutic strategy. Lastly, our unexpected finding of improved RNA packaging and luciferase induction by mutations within the N protein point to a previously unknown strategy for coronaviruses to evolve improved viral fitness. Although the mechanism for this improvement remains unclear, this finding is consistent with recent reports that the Delta variant (containing N:R203M) generates 1000-fold higher levels of RNA within patients (*26*). Our results point to a new and unanticipated mechanism that could explain why the SARS-CoV-2 Delta variant demonstrates improved viral fitness.

