## Supplemental Figures for "Rapid assessment of SARS-CoV-2 evolved variants using virus-like particles"

### Supplementary Figure 1

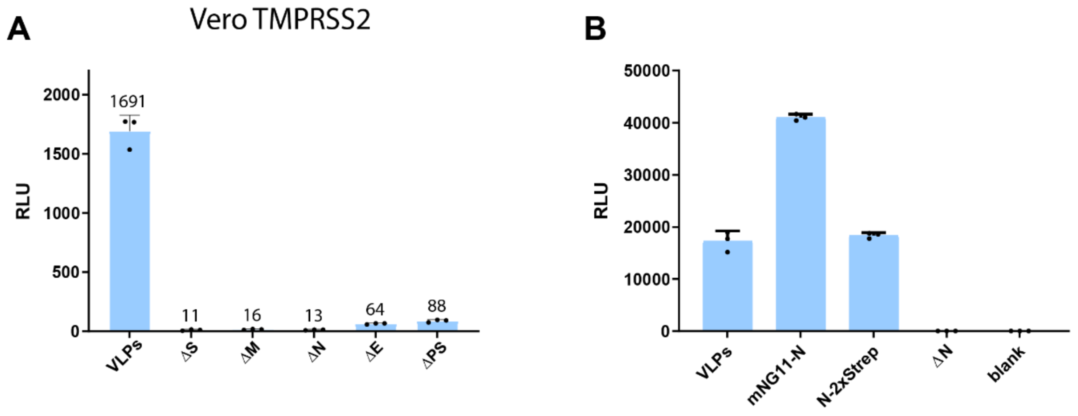


**Figure S1: Requirements for induced expression by SC2-VLPs.** A) Luminescence measured from Vero E6 cells incubated with supernatants containing SC2-VLPs as well as supernatants of cells missing either S, M, N, E or the packaging signal (PS). B) Luminescence from receiver cells after incubation with SC2-VLP containing supernatants as well as supernatants from cells transfected with N containing tags (mNG11-N: N with amino-terminal mNG11 tag and N-2xStrep: N with carboxy-terminal 2xStrep tag).

### Supplementary Figure 2

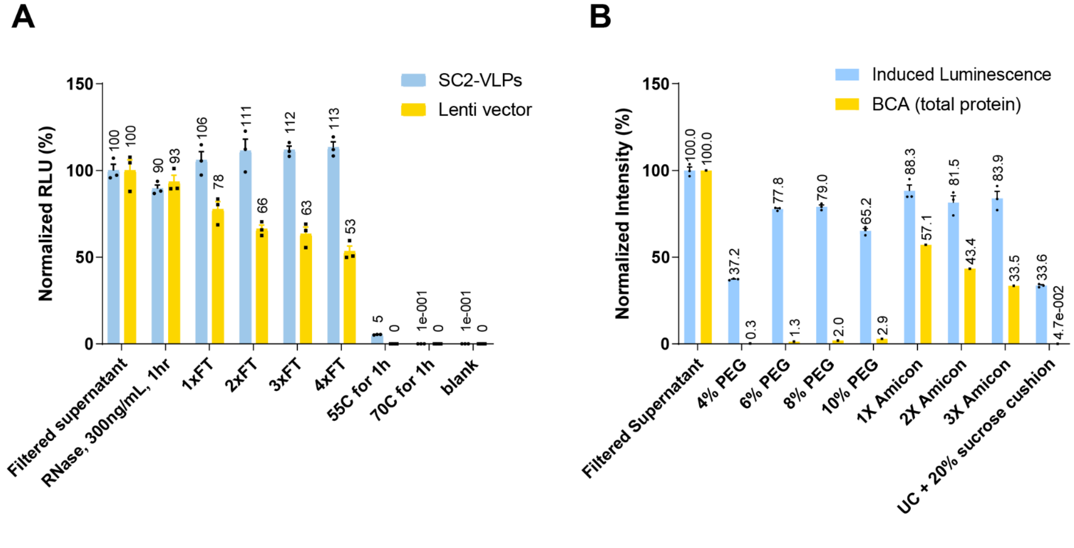


**Figure S2: Characterization of SC2-VLP stability and purification.** A) Luminescence induced in receiver cells from SC2-VLPs after treatment with ribonuclease or 1-4 cycles of freeze-thaw or incubation at 55°C and 70°C respectively. All values normalized to the original supernatant. Lentiviral particles encoding luciferase shown as comparison. B) Induced luminescence from SC2-VLPs purified/concentrated using different methods compared to total protein measurement from the same samples using bicinchoninic acid (BCA) assay.

### Supplementary Figure 3

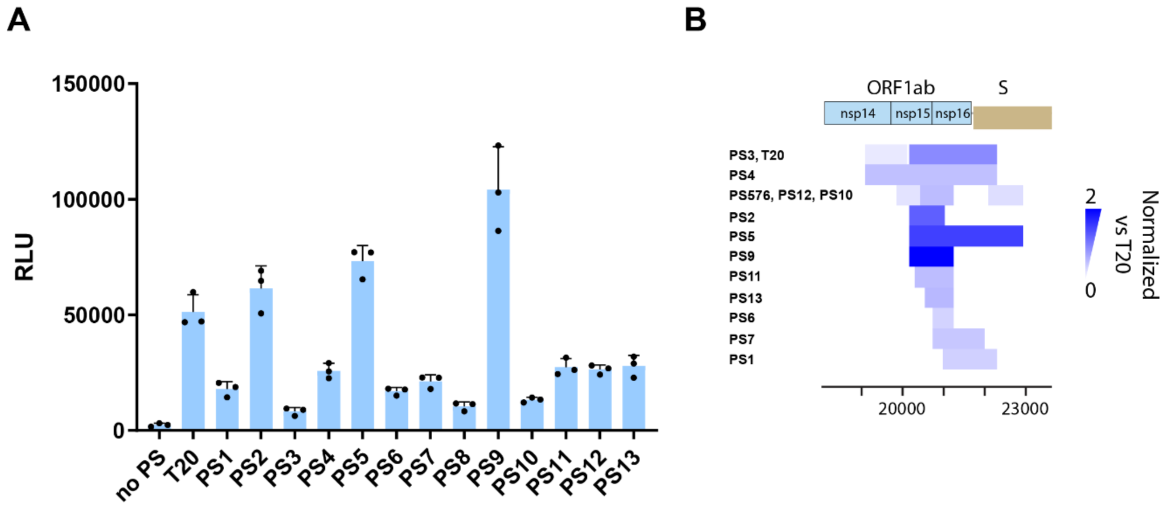


**Figure S3: Minimal sequence required for specific packaging into SC2-VLPs.** A) Induced luminescence in receiver cells after incubation with SC2-VLPs containing a transcript expressing luciferase. The luciferase transcript contains varying segments from SARS-CoV-2 shown graphically in (B). “no PS” indicates luciferase only transcript. Color in (B) indicates the observed luminescence normalized to the T20 transcript.

### Supplementary Figure 4

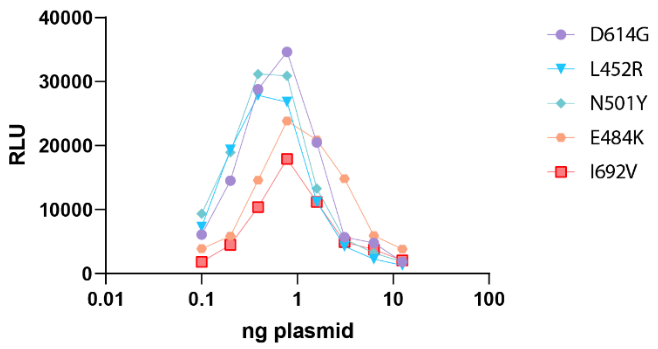


**Figure S4: Effect of spike mutations on SC2-VLP induced luminescence.** Induced luminescence from receiver cells incubated with SC2-VLPs containing varying concentrations and mutations within S plasmid. S plasmid ranging from 0.1 ng to 12.5 ng was added to each well of a 24-well plate. Total DNA used for transfection (N, M-IRES-E, T20) was 1 µg for each well.

### Supplementary Figure 5

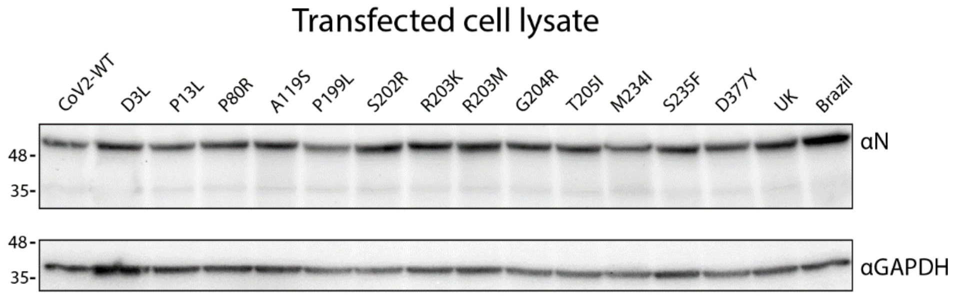


**Figure S5:** Expression levels of nucleocapsid containing mutations. Western blot of lysates from packaging cells transfected with N mutations stained using anti-N antibody (top) and anti-GAPDH antibody (bottom). Expression levels are similar between mutants and do not correlate with induced luminescence from SC2-VLPs made from these mutants.

**Materials and methods**

*Cloning for plasmids encoding structural proteins:* pcDNA3.1 backbone plasmids were generated encoding N, and M-IRES-E. Sequences for E, M and N were PCR amplified from codon optimized plasmids were gifts from Nevan Krogan (Addgene plasmid # 141385, 141386, 141391, ). pcDNA3.1-SARS2-Spike was a gift from Fang Li (Addgene plasmid # 145032). Site directed mutagenesis (NEB) was used to remove the C9-tag and introduce the D614G mutation.

*Cloning of SARS-CoV-2 genome tiled segments*: RNA was extracted from SARS-CoV-2 (Washington isolate) viral supernatant inactivated in Trizol by phase separation. RNA was reverse transcribed using protoscript II (NEB) and tiled segments (T1-T28) were PCR amplified from cDNA using primers compatible with ligation independent cloning (LIC). Tiles were cloned into a plasmid containing luciferase with a LIC destination site in the 3'UTR.

*SC2-VLP production:* For a 6-well, plasmids Cov2-N (0.67), CoV2-M-IRES-E (0.33), CoV-2-Spike (0.0016) and Luc-T20 (1.0) at indicate mass ratios for a total of 4 µg of DNA were diluted in 200 µL optimem. 12 µg PEI was diluted in 200 µL optimem and added to plasmid dilution quickly to complex the DNA. Transfection mixture was incubated for 20 minutes at room temperature and then added dropwise to 293T cells in 2 mL of DMEM containing fetal bovine serum and penicillin/streptomycin. Media was changed after 24 hours of transfection and At 48 hours post-transfection, VLP containing supernatant was collected and filtered using a 0.45 µm syringe filter. For other culture sizes, the mass of DNA used was 1 µg for 24-well, 4 µg for 6-well, 20 µg for 10-cm plate and 60 µg for 15-cm plate. Optimem volumes were 100 µL, 400 µL, 1mL and 3mL respectively and PEI was always used at 3:1 mass ratio.

*Luciferase readout*: In each well of a clear 96-well plate 50 µL of SC2-VLP containing supernatant was added to 50 µL of cell suspension containing 30 000 receiver cells (293T ACE2/TMPRSS2). Cells were allowed to attach and take up VLPs overnight. Next day, supernatant was removed and cells were rinsed with 1X PBS and lysed in 20 µL passive lysis buffer (Promega) for 15 minutes at room temperature with gentle rocking. Lysates were transferred to an opaque white 96-well plate and 50 μL of reconstituted luciferase assay buffer was added and mixed with each lysate. Luminescence was measured immediately after mixing using a TECAN plate reader.

*VLP purification using sucrose cushion*: SC2-VLP produced in 10-cm plates (10 mL of culture) were added to 13.2 mL ultracentrifuge tubes. 1 mL of 20% sucrose was underlaid using a 4" blunt needle. VLPs were centrifuged for 2 hours at 28 000 RPM using a SW41 Ti swinging bucket rotor. Supernatant was removed and ultracentrifuge tubes were inverted for 5 minutes on a paper towel with gentle tapping to remove remaining supernatant. VLPs were resuspended in 50 μL phosphate buffered saline for further experiments.

*SC2-VLP PEG precipitation*: 0.136 volumes of polyethylene glycol stock (50% PEG, 2.2% NaCl) was added to filtered supernatant containing SC2-VLPs to achieve a final concentration of 6% PEG. Solution was mixed thoroughly and precipitation was allowed to proceed for 2hrs at 4°C and then centrifuged at 2 000g for 20 minutes. Supernatant was discarded and VLPs were resuspended in PBS.

*SC2-VLP concentration using Amicon filters:* 0.5 mL filtered supernatant was added to 0.5 mL 100 kDa molecular weight cutoff amicon filters and centrifuged for 30 minutes at 2 000g. Concentrate was diluted in 1X PBS containing 0.02% tween 20 for all wash steps.

*Western blot cell lysate and VLPs*: For western blots of lysates, media was removed and cells were rinsed with PBS. Cells were then lysed for 20 minutes in RIPA lysis buffer containing Halt protease and phosphatase inhibitor cocktail. For western blots of ultracentrifuge concentrated VLPs, 10 mL of VLP supernatant from a 10-cm plate was pelleted (28000 RPM, 2hrs, SW41 Ti, 1mL 20% sucrose cushion), the supernatant was discarded and VLPs were resuspended in 50 μL of PBS. 15 μL of concentrated VLPs were used to western blot. Laemmli loading buffer (1x final) and dithiothreitol (DTT, 40 mM final) was added to lysates or VLP solution and heated for 95°C for 5 minutes to lyse VLPs and denature proteins. Samples were loaded on to 12-40% gradient gels (Biorad) and transferred to a PVDF membrane (Biorad). Membrane was blocked in 10% NFDM and stained with primary antibody: anti-N (abcam ab273434, 1:500 dilution), anti-S (abcam ab272504, 1:1000), anti-GAPDH (Santa Cruz sc-365062, 1:1000), antip24 (Sigma, 1:2000) for 2 hours at room temperature. Blots were rinsed with TBS-T three times for 10 minutes each and stained with secondary (mouse: abcam ab205719, or rabbit: invitrogen, 65-6120, 1:5000). Imaged using pierce chemiluminescence kit and Biorad Chemidoc imager.

*Sucrose gradient fractionation:* 10% to 40% sucrose gradient was prepared using a gradient mixer in 13.2 mL ultracentrifuge tubes. Concentrated and resuspended SC2-VLPs were overlaid on top of the gradient and centrifuged in a SW41 Ti rotor for 3 hours at 28 000 RPM. Gradient was fractionated from the bottom using a 4" blunt needle and a peristaltic pump. For cell infection, each fraction was diluted 20X and added to 293T cells expressing ACE2/TMPRSS2. Luciferase signal was measured the next day.

*GFP-VLPs and flow cytometry.* GFP was cloned into the luciferase destination vector (Luc-no PS) and Luc-PS9 to generate GFP-LIC and GFP-PS9. VLPs were generated in 10-cm plates and concentrated through a 20% sucrose cushion. 50 µL of concentrated VLPs were added to each well of a 24-well plate along with 120 000 receiver cells (293T ACE2/TMPRSS2). Cells were incubated with VLPs overnight and GFP expression was measured the next day using flow cytometry.

*Northern Blot:* VLPs collected from a 10-cm plate were concentrated by ultracentrifugation through a 20% sucrose cushion (28000 RPM, 2hrs, SW41 Ti). The supernatant was discarded and VLPs were resuspended in 50 µL of PBS. 20 µL of concentrated VLPs were used for Northern blotting. VLPs were lysed by adding 500 µL of Trizol (Sigma) and RNA was extracted by phase separation, precipitated with isopropanol with GlycoBlue and washed with 75% ethanol. RNA was resuspended in 30 µL of water, added to 30 µL 2x RNA Loading Dye (NEB) and denatured at 65°C for 15 minutes then loaded onto a 1% agarose gel containing 1X MOPS and 4% formaldehyde. Samples were run at room temperature for 12hrs at 20V and transferred by capillary action to Nylon membrane. The membrane was hybridized with a $^{32}P$-labeled luciferase DNA probe (Promega) and visualized using a phosphoscreen on a Typhoon imager (GE).
